# Simple One-step Vial-based Pretreatment for Deep Single-cell Proteomics and Its Application to Oocyte Aging

**DOI:** 10.1101/2024.12.03.626713

**Authors:** Hui Zhang, Hailu Zhang, Chuanxi Huang, Qing Zeng, Chunyan Tian, Fuchu He, Yun Yang

## Abstract

Single-cell proteomics is a pivotal technology for studying cellular phenotypes, offering unparalleled insights into cellular heterogeneity and dynamic functions. Technical improvement in mass spectrometry instrument and sample preparation has made single-cell proteomics feasible in recent years. Yet, developing a simple and robust sample preparation method to enable deep proteomics profiling of single cells remains a significant challenge. Herein, we developed a simple one-step vial-based pretreatment (SOViP) for deep label-free single-cell proteomics. SOViP integrates all sample preparation procedures into a single step in autosampler vials, yet it is highly efficient and high-throughput. SOViP can be finished within ∼2 h, with hands-on time limited to merely a few minutes. We show that on average over 6,500 can be quantified from a single mouse oocyte using SOViP. In total 6,983 protein groups were identified from single mouse oocytes across an entire reproductive lifespan, offering a valuable proteomics resource for oocyte aging. Unique molecular characteristics of oocytes at different ages were revealed, and a classifier consisting of ten proteins demonstrated accurate age-group classification and fertility-level prediction. Although demonstrated using mouse oocytes in this study, SOViP is adaptable to rare cell samples and other large cells, including follicles and preimplantation embryo cells, among others.

## INTRODUCTION

Single-cell analysis has become a key technology to elucidate cellular heterogeneity and dynamic functions.^1,2^ Techniques such as single-cell DNA sequencing (scDNA-seq) and single-cell RNA sequencing (scRNA-seq) are relatively mature and have been widely applied. However, genomics and transcriptomics provide only indirect insights into cellular states. Proteins are the workhorse for virtually all cellular processes, but the levels of protein and mRNA are only moderately correlated.^3,4^ Therefore, single-cell proteomics is an indispensable tool in elucidating the intricacies of cellular heterogeneity at the phenotype level. Unlike DNA and RNA whose signals can be amplified by robust polymerase chain reaction-based methods, single-cell proteomics is still technically challenging since no comparable amplification methods currently exist for proteins.

Due to advances in sample preparation and mass spectrometric instrumentation, a few studies have achieved proteomics profiling of single mouse oocytes or embryos in recent two years. Li et al. established an efficient and simplified single-cell proteomics (ES−SCP) workflow to realize proteomics profiling at the single-oocyte level.^5^ In ES−SCP workflow, oocytes were lysed through sonication and then heated for 15 min at 90 °C in a metal bath to denature proteins. After cooling to room temperature, proteins were digested at 37 °C overnight and desalted with a C18 stage tip. Li et al. quantified more than 4,000 protein groups from a pool of only 15 oocytes and more than 1,500 protein groups from single oocytes. Huang et al. compared aged and young mouse and human oocytes using multi-omics analysis.^6^ Oocytes were sonicated three times in a lysis buffer containing 8 M urea, and then centrifugated to remove remaining debris. Next, the collected supernatant was reduced, alkylated, digested by trypsin, and then desalted using a C18 SPE column. Approximately 1,100 protein groups were identified from single mouse oocytes, whereas nearly 10,000 genes were detected via scRNAseq. Very recently, Ye et al. developed One-Tip method, identifying ∼5,400 proteins from single zygotes by narrow-window data-independent acquisition (nDIA) analysis with the current state-of-the-art mass spectrometer Orbitrap Astral.^7^ The One-Tip sample preparation workflow consisted of eight steps, including multiple manual centrifugation steps. In addition, One-Tip method relied on specialized equipment and consumables such as Evosep and Evo-tips, which may limit its broader applicability.

In addition, single-cell multi-omics of mouse oocytes have been reported. Jiang et al. developed a single-cell simultaneous transcriptome and proteome (scSTAP) analysis platform based on microfluidics.^8^ Single oocytes were pre-lysed by Lys-C solution and then lysed by RapiGest. After the cell lysis, cell lyses were split into two aliquots-one for transcriptome analysis and the other for proteomic analysis. On average 2,663 protein groups and 19,948 genes were quantified from single mouse oocytes. He et al. developed an on-capillary alkylation microreactor (OCAM) to enable simultaneous proteome and metabolome profiling from the same cell.^7^ In OCAM, proteins were first covalently bound to an iodoacetic acid functionalized open-tubular capillary micro-reactor, and metabolites were eluted before on-column digestion of captured proteins. An average of 2,944, 3,039, and 3,085 proteins were quantified from single oocytes at GV (germinal vesicle), GVBD (germinal vesicle breakdown) and MII (metaphase II) stages, respectively, along with a total of 171 metabolites.

All the above-mentioned methods require laborious, multi-step operations for sample pretreatment, and a simple and robust method is still lacking to minimize variations introduced at each step. Herein, we reason that the routinely applied sonication, desalting and acidification steps can be omitted, and thus developed a simple one-step vial-based pretreatment (SOViP) for deep single-cell proteomics. In SOViP, we integrated all the sample preparation steps into a single incubation step to realize lysis and digestion simultaneously. The hands-on time is merely a few minutes for a single cell, and the samples are ready for nanoLC-MS/MS analysis immediately after incubation. To avoid cross-contamination and sample loss, we replaced the widely-used 384-well plates with autosampler vials. To ensure accessibility and low cost, we demonstrated that the incubation could be performed in a thermostat water bath. In addition, tens to hundreds of samples can be processed in a single batch, enabling SOViP a robust, simple and high-throughput method for broad applications.

Infertility is estimated to affect 8-12% of reproductive-aged couples worldwide, with declining fertility being one of the leading causes.^9,10^ Reduced oocyte quality due to aging is one of the most critical factors impacting female fertility. Single oocytes are heterogeneous and dynamic, however, proteomics studies at the single-cell level for oocyte aging are still scarce. The mechanisms underlying oocyte aging are still incompletely unraveled. To facilitate oocyte aging studies, we applied SOViP to profile mouse oocytes at different ages herein. On average, over 6,500 protein groups can be quantified from single oocytes. To the best of our knowledge, this study represents the first single-cell proteomics investigation of oocyte aging with deep coverage, providing a valuable resource for understanding oocyte quality and female fertility.

## EXPERIMENTAL SECTION

### Collection of single mouse oocytes

All animal protocols were carried out in accordance with the rules and guidelines of Medical Ethical Committee of the Beijing Institute of Lifeomics. Female mice of various ages, all on a C57BL/6 background, were raised under specific pathogen-free (SPF) conditions in individually ventilated cages.

To collect oocytes at Germinal vesicle (GV) stages, female mice at different ages were intraperitoneally injected with 5 IU pregnant mare serum gonadotrophin (PMSG). The intact ovaries were taken after 48 h, and their surfaces were then punctured with needles to release oocytes. Granule cells were carefully removed by gently pipetting oocytes repeatedly in pre-warmed M2 medium. Next, the oocytes were individually isolated and collected using a mouth pipette with an internal diameter slightly larger than the oocyte, under a stereoscopic microscope.

### Proteomics sample preparation

Single, clean oocytes collected from the above steps were transferred into 3 μL of master mix at the bottom of 384-well plates (Eppendorf, Protein LoBind, order No. 0030624300) or autosampler vials (Waters, QuanRecovery). The master mix, adapted from Ctortecka et al.,^11^ contained 0.2% n-Dodecyl-β-D-Maltoside (DDM), 100 mM N-2-hydroxyethylpiperazine-N-2-ethane sulfonic acid (HEPES) and 20 ng/μL Trypsin. After sealing with self-adhesive sealing films or screw caps, 384-well plates and autosampler vials were incubated at 37.0 °C in a commercial cellenONE system (85% humidity) or a thermostat water bath.

Human embryonic kidney (HEK) 293T cells were cultured in Dulbecco’s Modified Eagle Medium (DMEM) supplemented with 10% (v/v) fetal bovine serum (FBS) and Penicillin/Streptomycin (1:1000) at 37°C in a 5% CO_2_ atmosphere. Subsequently, 293T cells were detached using trypsin treatment and underwent thorough washing with phosphate buffered saline (PBS) three times. Then, lysis buffer was added and the lysis was assisted by non-contact sonication. The composition of the lysis buffer included 40 mM HEPES (pH 7.5), 300 mM NaCl, 1.2 M guanidine HCl, 2% (w/v) DDM, and a protease inhibitor mixture. Protein concentration was measured by nanodrop One. Different amounts of 293T cell lysates were added into 3 μL of master mix at the bottom of 384-well plates or autosampler vials and then incubated as described above.

All samples were kept at -80 °C prior to nanoLC-MS analysis. The sealing films of 384-well plates or the caps of vials were cleaned before injection. Note, no desalting or acidification steps were included in this protocol.

## Data availability

The mass spectrometry proteomics data related to this study have been deposited to iProX^12^ database (https://111.198.139.98/page/home.html) under the accession number IPX0010366000.

## RESULTS AND DISCUSSION

### Design of the SOViP strategy

In single-cell proteomics, it is well-known that reducing reactor volume minimizes contact surfaces, thereby decreasing sample loss due to non-specific binding to reactor surfaces.^13^ Consequently, successful single-cell proteomics for somatic cells are predominantly achieved using various microfluidic platforms with nanoliter-scale microreactors, such as oil-air-droplet (OAD) chip^14^, nanodroplet processing in one-pot for trace samples (nanoPOTS)^15,16^, sequential operation droplet array (SODA) system^17^, nested nanoPOTS (N2) chip^18^ and BOXmini SCP^19^. Alternatively, 1536-well plates^20^ and 384-well plates^21^ integrated with the cellenONE system have also been reported.

Oocytes and embryos are much larger than somatic cells, containing 100 times more protein amounts (∼20 ng for mouse oocytes and ∼100 ng for human oocytes) than somatic cells (∼200 pg). Though 384-well plates are compatible with direct injection into some liquid chromatography systems, the inadequate leakproofness of adhesive sealers and the remaining adhesive on sealers often cause problems (this will be detailed later). With the ultrahigh sensitivity of state-of-the-art MS instruments, such as timsTOF SCP, Orbitrap Astral and timsTOF Ultra 2, a 10-ng injection is sufficient to achieve good proteome coverage. We hypothesize that the slightly higher sample loss associated with larger reactors may not significantly compromise results in this context. Thus, autosampler vials with superior leakproofness are chosen as reactors for oocytes in this study. In addition, 384-well plates require incubation in a cellenONE system, which supplies controlled temperature and high-humidity water vapor generated from ultrapure water. Whereas for autosampler vials with robust leakproofness, routinely used thermostatic water bath may serve as their incubation chamber. Moreover, sample preparation should involve minimal steps to reduce time and variability.

To make it as simple and stable as possible, we developed the SOViP method, which integrated all the sample pretreatment steps into a single step using ready-to-inject autosampler vials (Fig. 1). The routinely required steps, including sonication, heating, cooling, reduction, alkylation, desalting, vacuum dryness, reconstitution and acidification, were omitted in our protocol. Instead, a single incubation step for simultaneous lysis and digestion remains in SOViP, ensuring the sample preparation is as simple as possible.

**Figure 1.**
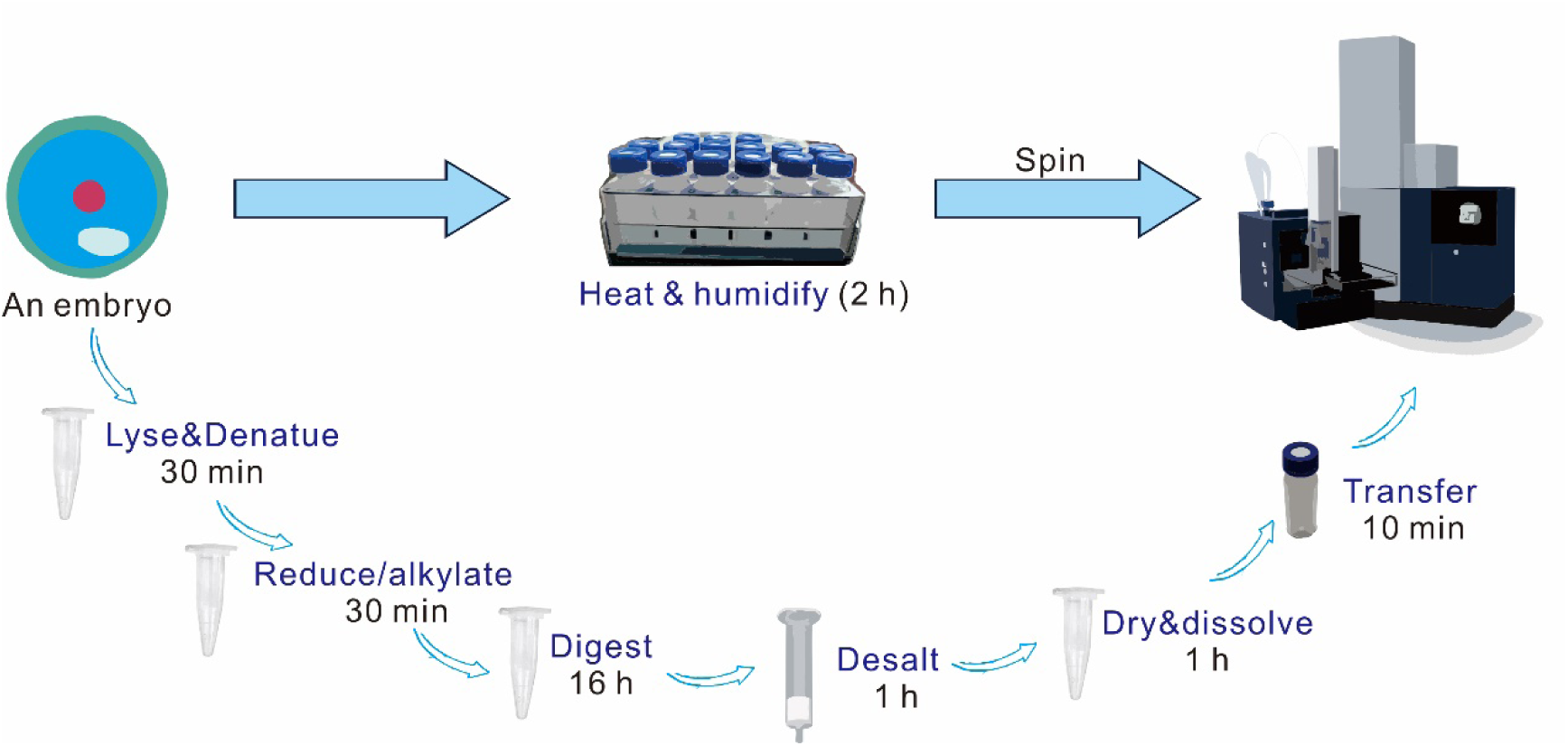
Scheme for the comparison of the simple one-step vial-based pretreatment (SOViP) method with a conventional multi-step method.

Consequently, the SOViP method can bring several merits:

1. Easy, fast and high throughput sample preparation: The only manual step involves placing single oocytes into the master mix at the bottom of autosampler vials, followed by a ∼2-hour incubation. Tens to hundreds of single cells can be processed per batch, with only a few minutes of hands-on time per cell.
2. Prevention of contamination and cross-contamination: Unlike 384-well plates or open-ended microfluidic chips, autosampler vails can be tightly sealed with screw caps, ensuring excellent leakproofness.
3. Improved injection completeness: Autosampler vials are better suited for injection needles than most 384-well plates, where incomplete injection is common.
4. ∼100% success rates with reduced variability: The adhesive from adhesive sealers can easily block injection needles/injection ports/plungers, resulting in incomplete injections or even failures. Meanwhile, the adhesive also increases the frequency of LC system maintenance, prolonging instrument downtime.
5. Simpler operation: When using 384-well plates, unused wells must be sealed and resealed multiple times during preparation, adding complexity. Autosampler vials eliminate this need.
6. Flexibility to incorporate sonication: Non-contact sonication can be employed to enhance lysis for certain sample types.
7. Extended sample storage: Samples in 384-well plates must be injected within ∼2 days due to evaporation, while those in autosampler vials remain stable for ∼2 weeks.
8. Cost efficiency: The cellenONE system is expensive and accessible in only a few laboratories, whereas most laboratories already have a thermostatic water bath.

### Evaluation of SOViP for cell lysate samples

We first compared the performance of 384-well plates vs. autosampler vials with 200 pg and 1 ng of 293T cell lysate (Figs. 2a-2b), amounts close to the protein amount in a single somatic cell. Out of the total ten tested samples, three failed in the 384-well plate group, whereas all samples yielded stable results in the autosampler vial group. This highlighted the superior stability afforded by using autosampler vials over 384-well plates. Nevertheless, excluding the failed injections (three data with less than 200 protein groups identified), 21% and 11% more protein groups were identified using 384-well plates compared to autosampler vials for 200 pg (2,168 vs. 1,799) and 1 ng (2,554 vs. 2,297) of 293T cell lysate, respectively; Meanwhile, 384-well plates led to 30% and 20% more peptide groups identified for 200 pg (10,056 vs. 7,727) and 1 ng (14,880 vs. 12,290) of 293T cell lysate, respectively. This is rational, as the reactor volume of 384-well plates is smaller than that of autosampler vials, leading to less sample loss caused by non-specific binding. Consequently, 200 pg benefit more than 1 ng from smaller reactor volume in our results.

**Figure 2.**
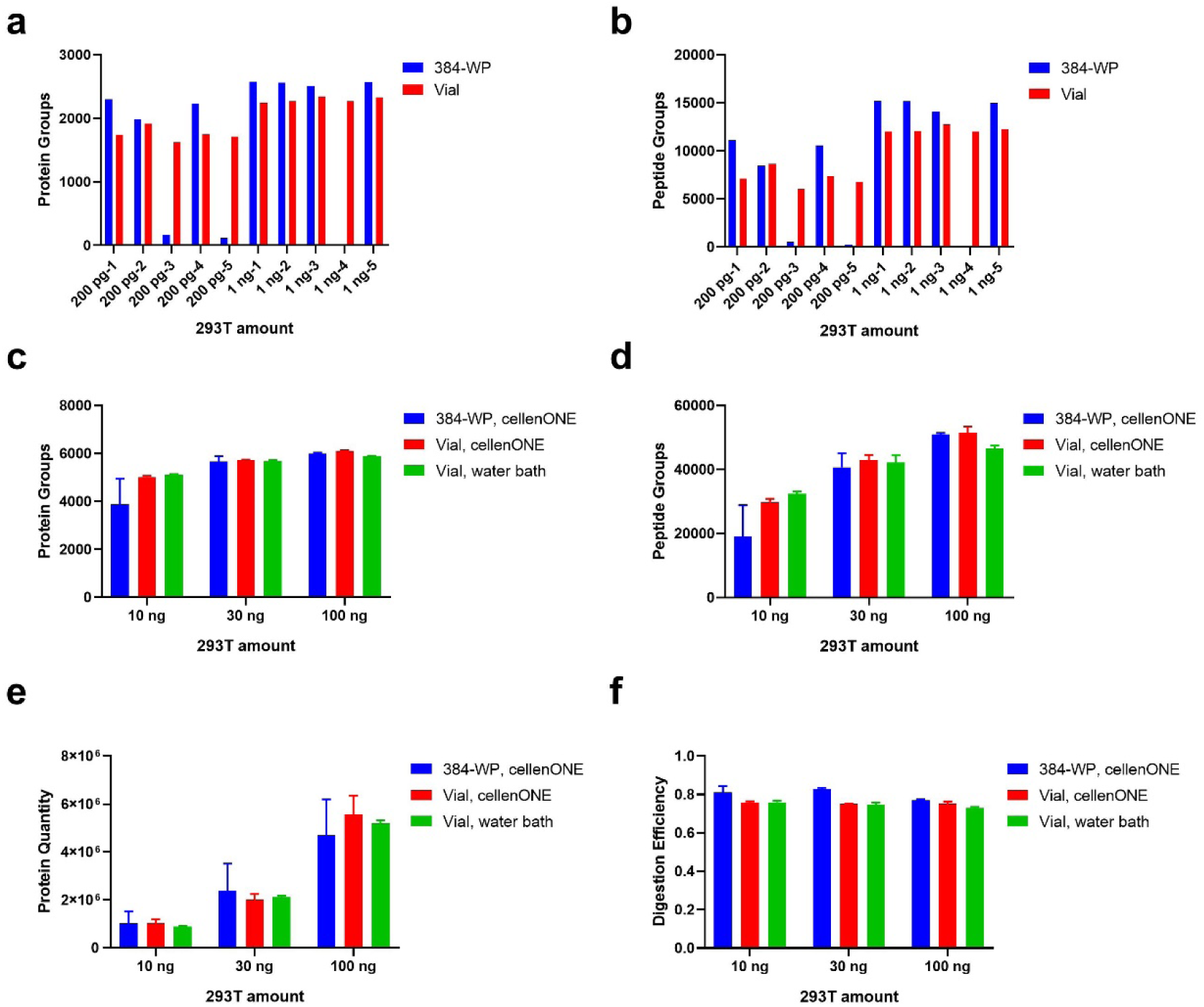
Comparison of 384-well plate and autosampler vial for processing different amounts of 293T cell lysate. **a-b** The number of protein groups (a) and peptide groups (b) identified from 200 pg and 1 ng of 293T cell lysate using a 384-well plate or vials. **c-d** The number of protein groups (c) and peptide groups (d) identified from 10, 30, and 100 ng of 293T cell lysate using a 384-well plate or vials in a cellenONE or a water bath. **e-f** Comparison of protein quantity (e) and digestion efficiency (f) for 10, 30, and 100 ng of 293T cell lysate processed using a 384-well plate or vials in a cellenONE or a water bath. “384-WP” and “Vial” indicate 384-well plate and autosampler vial, respectively.

Next, we compared these two reactors for higher protein amounts and obtained different outcomes. 10, 30, and 100 ng of 293T cell lysate span the protein range of mouse to human oocytes at various stages. As shown in Figs. 2c-2d, the 384-well plate group still showed significantly higher variability than the autosampler vial group in terms of protein groups, peptide groups, protein quantity and digestion efficiency. For 10-100 ng of 293T cell lysate, more protein groups and peptide groups were identified using autosampler vials, in contrast to the results for 200 pg and 1 ng. This indicates that sample loss due to non-specific binding is minimal for high protein samples, while sample loss due to evaporation during incubation is more significant when sealing is inadequate. This suggests that sample loss due to non-specific binding is minimal for high protein samples, while sample loss due to evaporation during incubation is more significant when sealing is inadequate. When using autosampler vials, 5,014 ± 49 protein groups were identified from 10 ng of 293T cell lysate, and this number increased to 5,714 ± 29 and 6113 ± 14 for 30 and 100 ng of 293T cell lysate, respectively. Not surprising, variation decreased along with increasing protein amount. In addition, digestion efficiency, represented by the relative percentage of fully tryptic peptides to all identified peptides, achieved close to 80% (Fig. 2f), which is satisfactory for a simple one-step and one-pot sample pretreatment.

Lastly, to enhance accessibility, we tested using a thermostatic water bath to replace cellenONE when using autosampler vials. A thermostatic water bath can supply stable temperature and high humidity, although not as precisely controlled as cellenONE. Moreover, the water vapor in thermostatic water is not clean, requiring that the reactor be well sealed. Encouragingly, the water bath achieved results similar to those of cellenONE, with no significant differences in protein groups, peptide groups, protein quantity, or digestion efficiency between the water bath group and the cellenONE group.

### Development of SOViP for mouse oocytes

The SOViP protocol for mouse oocytes is depicted in Fig. 3a. Female mice were intraperitoneally injected with PMSG to accelerate follicular development. Granule cells were carefully removed, and single oocytes were isolated and collected under a stereoscopic microscope. Single oocytes were directly transferred to the bottom of autosampler vials using mouth pipette. After a 2-hour incubation, the samples in the autosampler vials were directly injected. The workflow is simple and straightforward (Fig. 3a).

**Figure 3.**
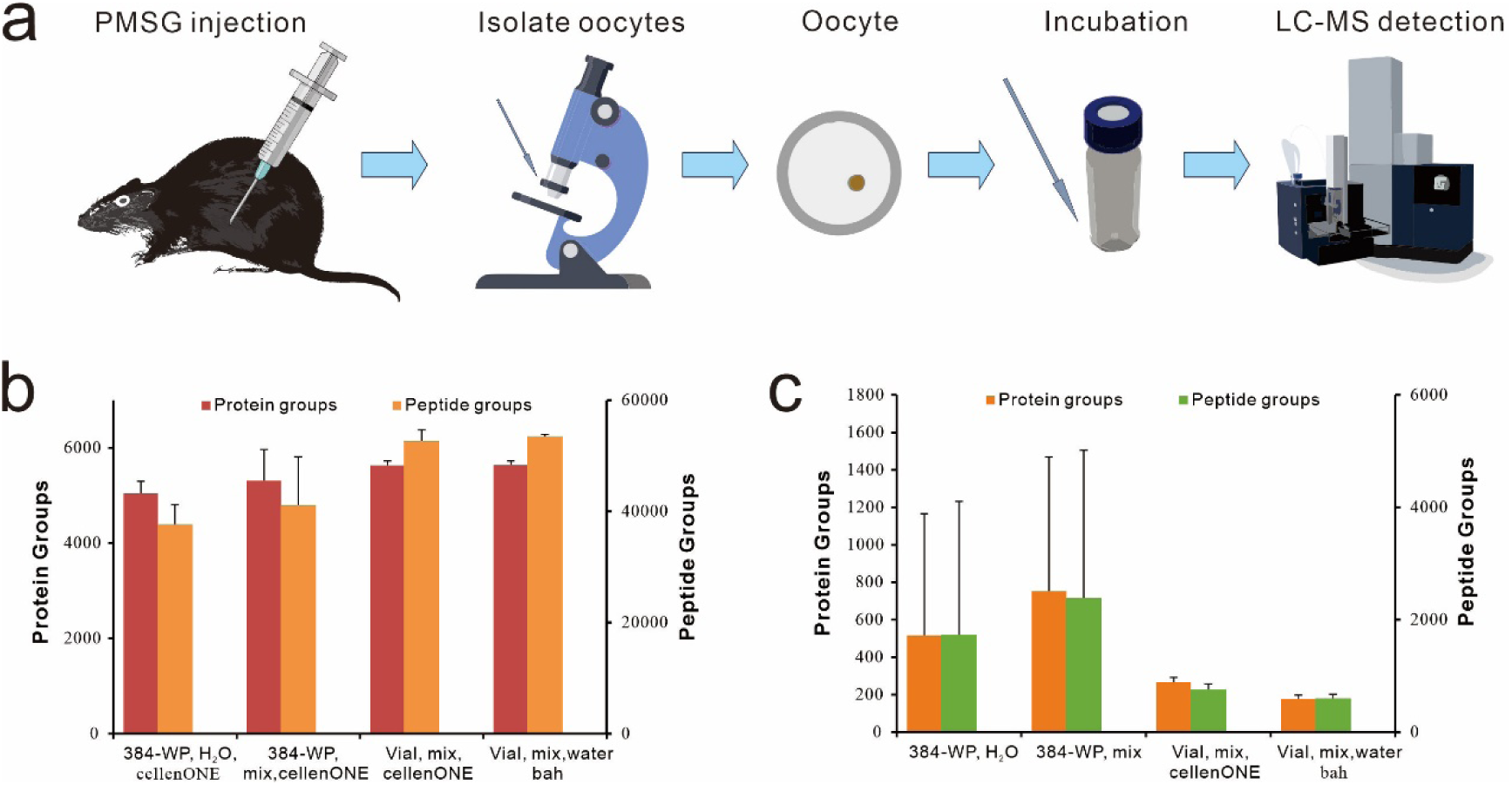
Method development for processing single mouse oocytes. **a** Workflow for processing single mouse oocytes using SOViP. **b** Comparison of the number of protein groups and peptide groups identified from single mouse oocytes using four protocols: Single mouse oocytes were kept in H_2_O, transferred to a 384-well plate, freeze-dried, and incubated with 3 μL of master mix in cellenONE (“384-WP, H_2_O” group); or directly transferred into 3 μL of master mix in a 384-well plate (“384-WP, mix” group). Alternatively, single mouse oocytes were transferred into 3 μL of master mix in vials and then incubated in either cellenONE (“Vial, mix, cellenONE” group) or a water bath (“Vial, mix, water bath” group). **c** Comparison of the number of protein groups and peptide groups identified from blank controls using the above four protocols.

During workflow optimization, we compared four protocols for single mouse oocytes: Oocytes were kept in H_2_O, transferred to a 384-well plate, freeze-dried, and then incubated in cellenONE after adding master mix (“384-WP, H_2_O” group); (2) Oocytes were directly transferred into master mix in a 384-well plate, and then incubated in cellenONE (“384-WP, mix” group); (3) Oocytes were directly transferred into master mix in autosampler vials and then incubated in cellenONE (“Vial, mix, cellenONE” group); (4) Oocytes were directly transferred into master mix in autosampler vials and then incubated in a water bath (“Vial, mix, water bath” group). The “384-WP, H_2_O” group involved additional transfer and freeze-drying steps but yielded lowest numbers of identified protein and peptide groups (Fig. 3b). Direct transfer into 384-well plates without freeze-drying yielded slightly higher protein and peptide group counts. (Fig. 3b). In contrast, the two autosampler vial groups, whether using cellenONE or a water bath, produced significantly higher protein and peptide group counts than either of the 384-well plate groups (Fig. 3b). Moreover, the autosampler vial groups exhibited much lower variability compared to the 384-well plate groups, consistent with the trend observed for cell lysates (Fig. 2).

To assess background contamination, we compared blank controls for those four protocols as well (Fig. 3c). When using 384-well plates, approximately 600 protein groups with large variations were detected. In contrast, merely ∼ 200 protein groups were detected in the two autosampler vial groups, likely originating from reagent and environmental background noise^8^. Overall, the autosampler vial-based protocols demonstrated clear advantages over the 384-well plate-based protocols for pretreating mouse oocytes.

### Application of SOViP in oocyte aging

Aging exerts multifaceted effects on the female reproductive system, with a notable decline in oocyte quality and an impediment to embryo developmental potential.^9^ In 2023, Isola et al. built a single-cell RNA sequencing atlas of the aging mouse ovary by comparing ovarian tissues from 3-month-old and 9-month-old mice.^22^ This study provided a comprehensive cellular map of the transcriptomic changes in the aging mouse ovary, revealing a doubling of immune cell populations in aged ovaries. Huang et al. performed single-cell proteomics, translatomics and transcriptomics analysis on GV-stage oocytes from aged and young mice and humans.^6^ Huang et al. identified a crucial m6A-containing YTHDF3-binding target Hells, and highlighted cross-species conservations and differences between aged mouse and human oocytes. Despite significant strides in research, the intricate mechanisms governing oocyte senescence remain incompletely elucidated, and effective approaches to improve oocyte quality are still lacking.^23^

Naturally-aged female mice are widely used as in vivo models for oocyte aging studies, because they exhibit physiological changes that resemble those of menopausal women, including irregular oestrus cycles. C57BL/6 mice, commonly used for such studies, are weaned at 3-4 weeks after birth and attain sexual maturity at 8-12 weeks of age.^24^ Studies have shown that the fertility of C57BL/6 female mice begins to decline from the highest level at approximately 6 months of age and drops markedly by 12 months.^23^ By 15 months, fertility is nearly exhausted^24^, when ovulated oocyte numbers largely diminish and oocyte quality decays. To comprehensively study reproductive senescence, we selected five age groups of wild-type female mice: 3 weeks (post-weaning), 8 weeks (adult), 6 months (mid-life), 12 months (late reproductive life), and 15 months (end of fertility).

Using the developed SOViP workflow, we conducted single-cell proteomics analysis to deeply profile the proteomes of mouse oocytes at the GV stage and to investigate reproductive aging at the single-cell level. We achieved deep proteome profiling of single mouse oocytes from weaning to fertility end period, using both cellenONE and water bath (Fig. 4, Table S2). Results from the two methods were highly similar, with cellenONE yielding only slightly higher numbers of identified protein and peptide groups (Figs. 4a–4b), consistent with previous findings (Fig. 3). Principal component analysis (PCA) and Pearson correlation coefficients demonstrated strong concordance between biological replicates and high correlation between the two preparation methods (Fig. S1a). Across the five age groups, proteomic analysis revealed dynamic patterns. On average, 6,009 protein groups were identified from single oocytes at 3 weeks, increasing significantly to over 6,500 in the 8-week to 12-month groups before declining sharply to ∼5,300 at 15 months, corresponding to the poor quality of oocytes at the end of fertility (Fig. 4c). A similar trend was observed for peptide group counts. This rise-and-fall pattern suggests that the expression of approximately 500 fertility-related genes is low at 3 weeks and that over 1,000 genes are downregulated at 15 months (Figs. 4c&4e). Noteworthily, the total protein quantity increased steadily from 3 weeks to 6 months but declined from 6 months to 15 months (Fig. 4d), indicating age-dependent differences in protein expression patterns. A high proportion of protein groups were shared between adjacent age groups (Fig. 4e), with 6,139 protein groups co-identified across all five age groups (Fig. S2), indicating most proteins continuously expressed from 3 weeks to 15 months. Remarkably, using the easy-to-use one-step SOViP protocol, we achieved dynamic ranges of quantified protein groups spanning six orders of magnitude (Fig. S3), capturing a high ratio of low-abundance proteins. Among them, 14 of the 36 reported pathogenic genes involved in human oocyte and early embryo development^25^ were quantified and marked in Fig. 4f. Additionally, we quantified 191 mouse transcription factors across a wide dynamic range (Fig. 4g, Table S1).

**Figure 4.**
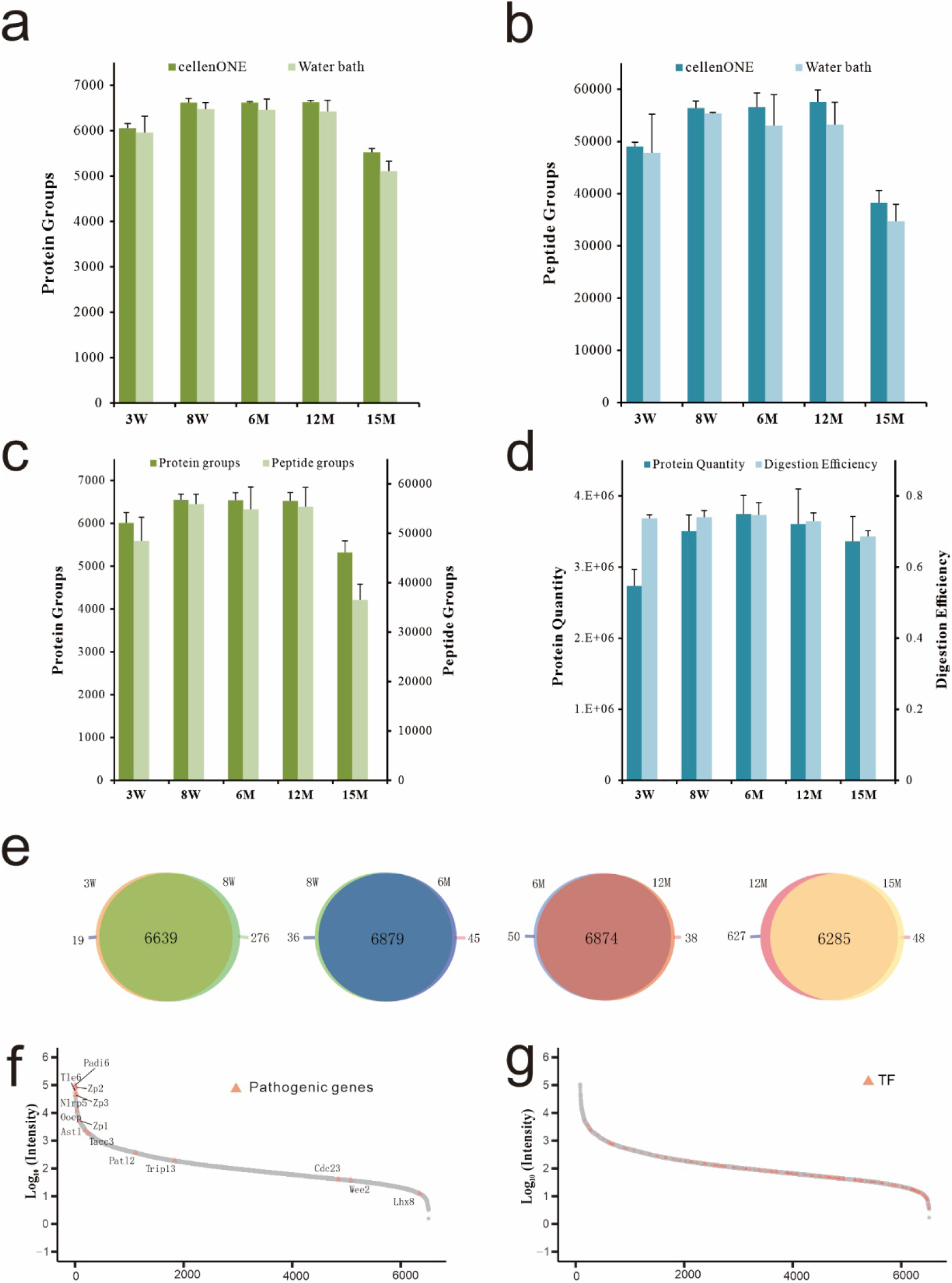
Application of SOViP to oocyte aging. **a-b** Comparison of the number of protein groups (a) and peptide groups (b) identified from single mouse oocytes at five ages incubated in cellenONE or a water bath. **c** The number of protein groups and peptide groups identified from single mouse oocytes at five ages. **d** Comparison of protein quantity and digestion efficiency for single mouse oocytes at five ages. **e** Venn diagrams showing the overlap of identified protein groups across four adjacent age groups. **f-g** Assessment of dynamic range based on protein abundance rank (mean values for the five age groups) and annotation of 198 mouse transcriptional factors (f) and 21 reported pathogenic genes involved in human oocyte and early embryo development (g). 3W, 8W, 6M, 12M, and 15M represent 3-week, 8-week, 6-month, 12-month, and 15-month samples, respectively.

Overall, a total of 6,983 protein groups were identified and 6,927 protein groups were quantified from single mouse oocytes (Table S2). On average, ∼6,500 protein groups were quantified from single oocytes aged 8 weeks to 12 months. Notably, detecting over 5,500 protein groups from a single mouse oocyte has not been reported previously, even using more advanced MS like Orbitrap Astral or timsTOF Ultra 2 combined with multi-step protocols.^26^ Therefore, our simple, one-step method represents a significant technical advancement in the pretreatment of single oocytes. Moreover, the five age groups from 3 weeks to 15 months provide a comprehensive single-cell proteomic resource for age-related fertility research.

### Oocyte characteristics in different ages

To investigate biological characteristics for oocytes in different ages from single-cell proteomics data, we then performed unsupervised hierarchical clustering of differentially expressed proteins. Data from the cellenONE group and the water bath group were combined for biological analysis, and comparisons between each pair of adjacent age groups are presented in Fig. 5. All data were correctly clustered in corresponding age groups, with representative biological processes (BPs) for each cluster highlighted in the boxes on the right. Gene ontology (GO) enrichment analysis of clusters revealed unique characteristics of these age groups. DNA methylation involved in gamete generation, DNA methylation, DNA modification or demethylation and DNA modification were enriched in 3-week oocytes, suggesting that 3-week oocytes in young mice immediately after weaning tend to complete essential developmental milestones and maintain viability. Regulations of lipid and lipoprotein, including lipid storage, protein lipid complex subunit organization, regulation of lipid localization, high density lipoprotein particle remodelling, positive regulation of lipid localization, regulation of plasma lipoprotein particle levels, were enriched in 8-week oocytes, indicating that oocytes extensively consumed lipids and lipoproteins during the process of sexual maturation. When compared with 8-week oocytes, upregulation of proteins in 6-month oocytes mainly related to mRNA processing, RNA splicing, RNA splicing via transesterification reactions, organic acid catabolic process, fatty acid catabolic process, small molecule catabolic process and monocarboxylic acid catabolic process, which shows the augmented RNA expression and catabolic processes in 6-month oocytes correlates with enhanced reproductive capability. For oocytes collected from 12-month-old mice, the upregulation of cysteine-type endopeptidase activity suggested an increase in apoptosis.^27^ Additionally, the upregulation of proteins primarily associated with the migration of mononuclear cells and myeloid leukocytes resulted from age-related senescence.^28,29^ The elevated responses to metal ions and the negative regulation of T-cell receptor and antigen receptor-mediated signalling pathways correlated with significant reproductive senescence. Meanwhile, the downregulation of fatty acid biosynthesis, lipoprotein lipase activity, and plasma lipoprotein particle clearance indicated abnormal lipid metabolism in oocyte senescence.^30,31^ Oocytes collected from 12-month-old mice exhibited higher ATP activity and electron transport, compared to 15-month-old oocytes which were at the end of fertility. As mitochondria are the primary organelles responsible for ATP production to support cellular biological processes, previous studies have shown that both the number and activity of mitochondria decrease with advancing age^32^. Electrons are transported to the mitochondrial membrane and participate in oxidative phosphorylation, while the mitochondrial membrane potential of oocytes increases with age.^33,34^ Reactive oxygen species (ROS), a byproduct of mitochondrial oxidative phosphorylation^35^, have been widely linked to age-related infertility in females, as studies indicate that increased ROS levels correlate with advancing age^36,37^. Together, the observed differences in ATP synthesis and electron transport between 12-month-old and 15-month-old oocytes highlight the progressive impairment of mitochondrial function with age. In 15-month-old oocytes, negative regulation of mammalian target of rapamycin (mTOR) signalling was elevated, which need further investigation.

**Figure 5.**
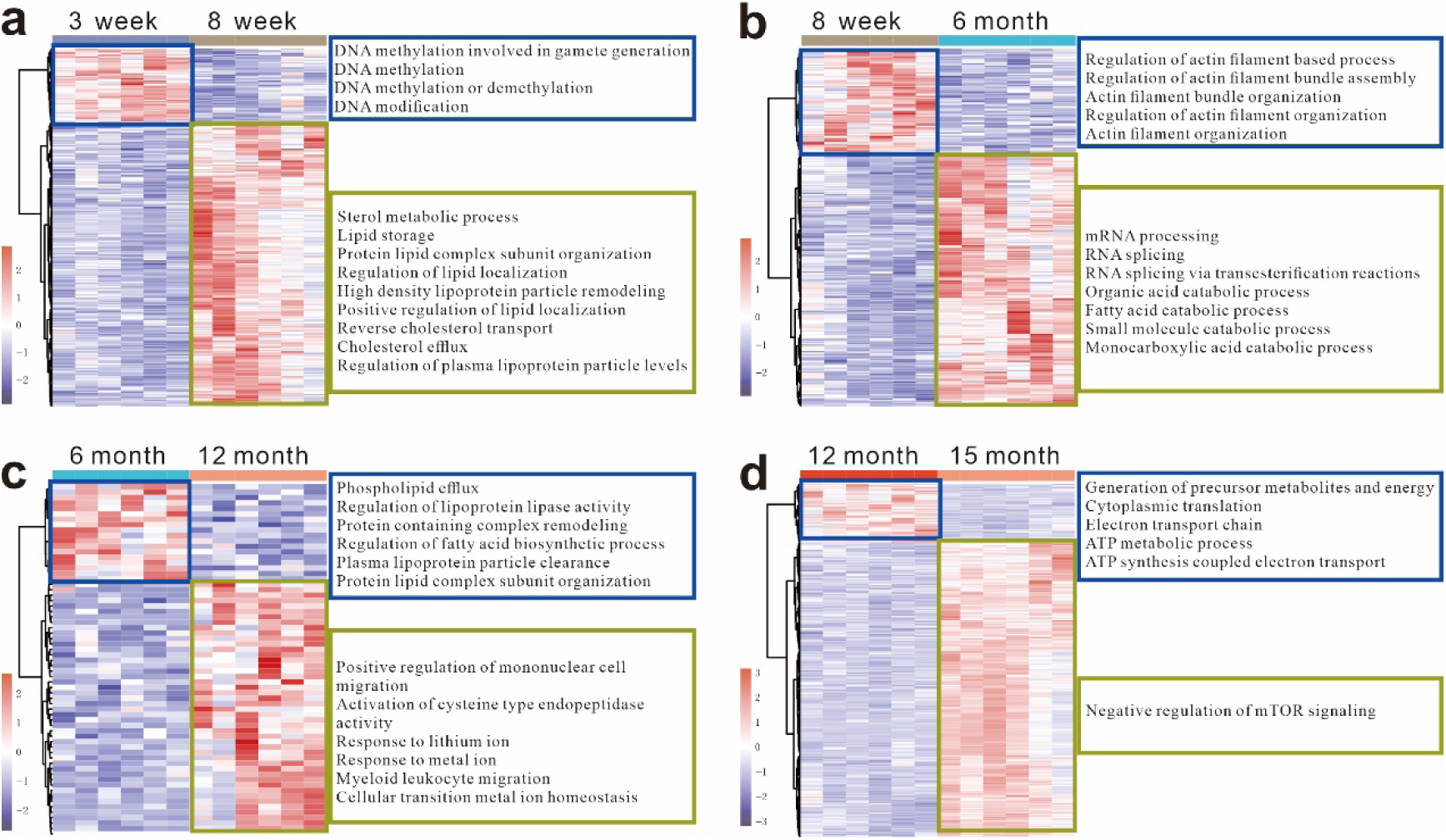
Oocyte characteristics across different ages. **a-d** Left: Unsupervised hierarchical clustering of proteins showing significant regulation (p < 0.05, one-sided ANOVA) and FC > 1.5 fold in each adjacent age groups: 3 weeks vs. 8 weeks (a), 8 weeks vs. 6 months (b), 6 months vs. 12 months (c), and 12 months vs. 15 months (d). Right: Gene ontology biological processes enrichment analysis of the protein clusters from the heatmap. Number of biological replicates: n=6 for all age groups.

### Biological insights into oocyte aging

To identify the significantly enriched functional or regulatory features governing mouse oocyte aging across all the five age groups, we grouped all the significantly regulated 3,203 protein groups into six clusters according to their expression patterns by means of Mfuzz algorithm (Figs. 6a-6b). Cluster 6 proteins (n = 135) showed a consecutively increasing trend. Cluster 6 proteins were enriched in mRNA and splicing activities, including alternative mRNA splicing via spliceosome, mRNA processing RNA splicing, regulation of mRNA processing, RNA splicing via transesterification reactions, regulation of mRNA splicing via spliceosome, regulation of mRNA metabolic process. In contrast, Cluster 4 proteins (n = 321) displayed trends opposite to those of Cluster 6 proteins and were enriched in organization of organelle organization, chromosome organization, and associated protein localization activities. Cluster 2 and Cluster 5 displayed similar rise-and-fall trends, with protein expression levels in both clusters decreasing sharply at 15 months. These proteins were primarily associated with energy consumption-related GO categories. Proteins in Cluster 3 (n = 282) exhibited a gentle increase at 8 weeks, followed by a gradual decline from 8 weeks to 12 months, and a sharp surge at 15 months. However, no significant BPs were identified for Cluster 3 proteins. Collectively, these findings underscore the close links between age-related protein abundance dynamics and distinct BPs.

**Figure 6.**
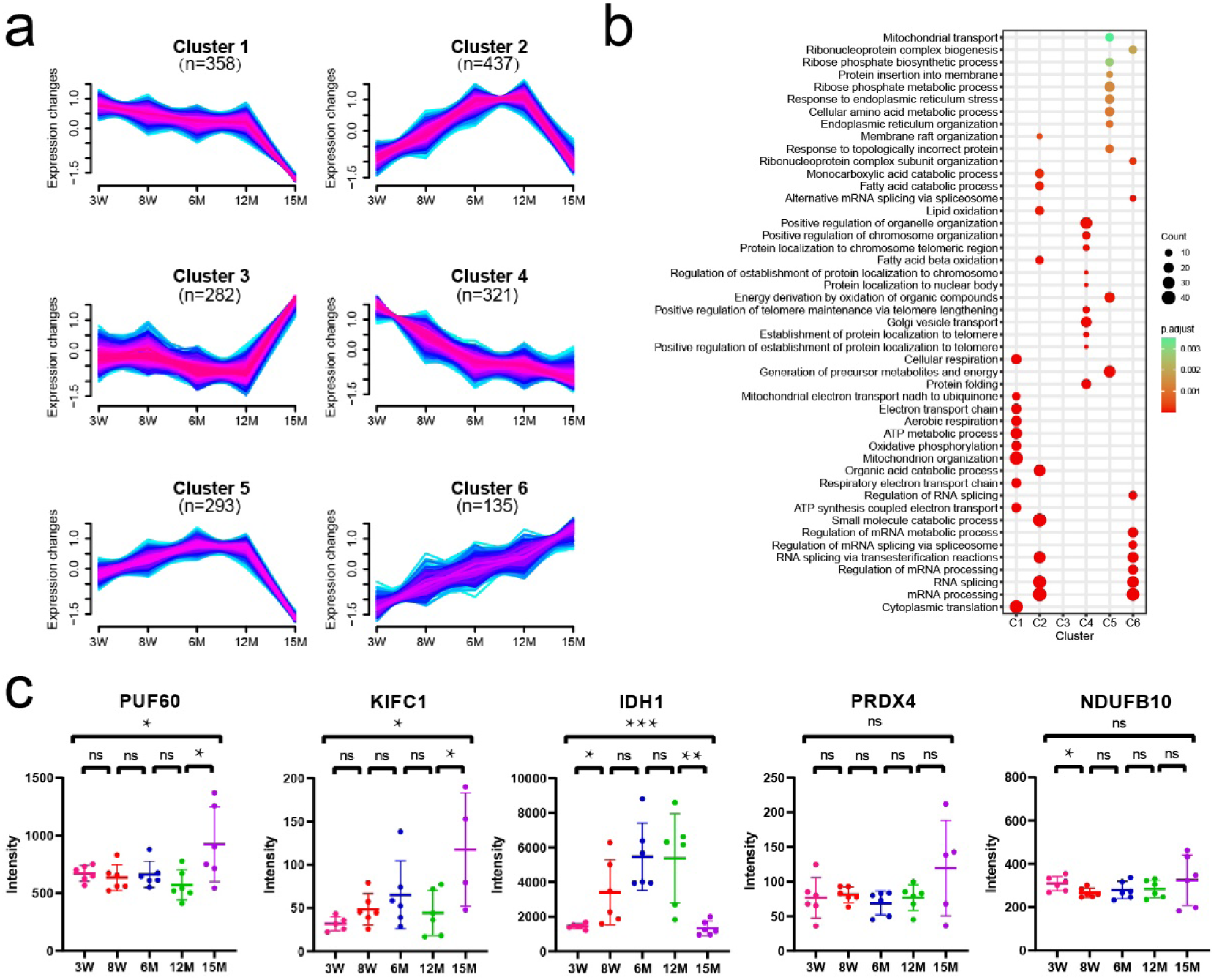
Biological insights of mouse oocyte aging. **a** Dynamic proteins were categorized into six clusters using Mfuzz soft clustering. A total of 3,203 dynamic proteins filtered by an ANOVA p-value < 0.05 were analyzed. Proteins were considered significant regulation across five groups if they presented a Benjamini– Hochberg (B–H) adjusted p-value of less than 0.05 from one-sided ANOVA. **b**. Representative biological processes from gene ontology (GO) enrichment analysis for each cluster are displayed. Dot size indicates the number of enriched proteins for each GO term, and the colour scale represents the statistical significance of each enrichment. **c.** Relative expression of five reported marker proteins related to fertility at different ages (n = 6). Data are presented as mean ± SD. Analyses were conducted using a two-tailed unpaired Student’s t-test (***p < 0.001, **p < 0.01, *p < 0.05, ns: not significant); 3W, 8W, 6M, 12M, and 15M indicate 3-week, 8-week, 6-month, 12-month, and 15-month samples, respectively.

Next, we investigated the abundances of five previously reported protein markers related to aging and reproductive capacity across the five age groups (Fig. 6c), since single-cell proteomics provides novel insights compared to bulk proteomics and single-cell RNA sequencing. PUF60 (Poly (U) Binding Splicing Factor 60) was reported as a potential core splicing factor responsible for the decline of oocyte developmental potential in reproductively aging mice, based on the observation that expression level of PUF60 progressively decreased with oocyte aging using tandem mass tag (TMT)-labelled bulk proteomics involving hundreds of oocytes.^23^ However, in our single-cell proteomics data, PUF60 remained stable from 3 weeks to 12 months before increasing at 15 months, suggesting that further research is required to elucidate its role in oocyte aging. Molecular motor KIFC1 (kinesin superfamily protein C1) was reported to be deficient in human oocytes, leading to spindle instability^38^. Similarly, KIFC1 depletion induced spindle instability in mouse oocytes. Nevertheless, in our data, KIFC1 levels did not decline with oocyte aging (Fig. 6c), indicating that KIFC1 may not significantly contribute to the decline of reproductive capacity in mice. Additionally, three antioxidant genes (IDH1, PRDX4 and NDUFB10) have been reported to negatively correlate with age in human granulosa cells.^39^ In our single-cell proteomics data of mouse oocytes, however, only IDH1 was significantly downregulated in low-fertility ages (low at 3-week and undetected at 15-month), indicating potential conflicting expression levels between oocytes and granulosa cells. Notably, substantial variability was observed within the same age group, reflecting cellular heterogeneity and underscoring the necessity and value of single-cell studies. Overall, single-cell proteomic data can provide novel biologically meaningful insights into the mechanisms governing oocyte aging. Yet, our findings require further investigation and validation in future studies.

### Machine learning-based classification of fertility ability

Assessing oocyte quality using biomarkers is important in assisted reproductive technology. To advance this, we developed machine learning-based models to classify and differentiate the five age groups and their corresponding fertility levels using our single-cell proteomics data. Because the sample size of this proof-of-concept study is limited, to alleviate overfitting, preliminary filtering of features by ANOVA was performed, and then a hybrid machine learning-based model with cross-validation were trained. Top 100 differentially expressed proteins (DEPs) were selected by ANOVA from the all 3,203 dynamic proteins, and hybrid SVM-RF models were trained to generate best classifiers to differentiate high (8W, 6M) versus low (3W, 12M, 15M) fertility ability. The best classifier consisted of ten proteins, Cpsf6, Aldh16a1, Dctn3, Snx27, Rpl10, Cbx1, Oat, Atp5f1d, Fdx1 and Sra1, and achieved a high AUC value of 1.000 in differentiating high versus low fertility ability (Fig. 7a). Importantly, those ten proteins can further accurately classify each of those five age groups and corresponding different fertility abilities as well. As shown in Figs. 7b-7f, the prediction accuracy achieved 1.000, 0.993, 0.938, 0.882 and 1.000 when classifying 3-week, 8-week, 6-month, 12-month and 15-month against the rest groups, respectively. Protein expression levels and prediction probability of those ten proteins for five age groups were depicted in Fig. S4. It is reasonable that the AUC value of 12-month is lower than the others, since the fertility difference of 12-month is smaller than that of 3-week and 15-month. Our research laid a foundation for discovering age-specific biomarker panels associated with different fertility levels in future large-scale studies.

**Figure 7.**
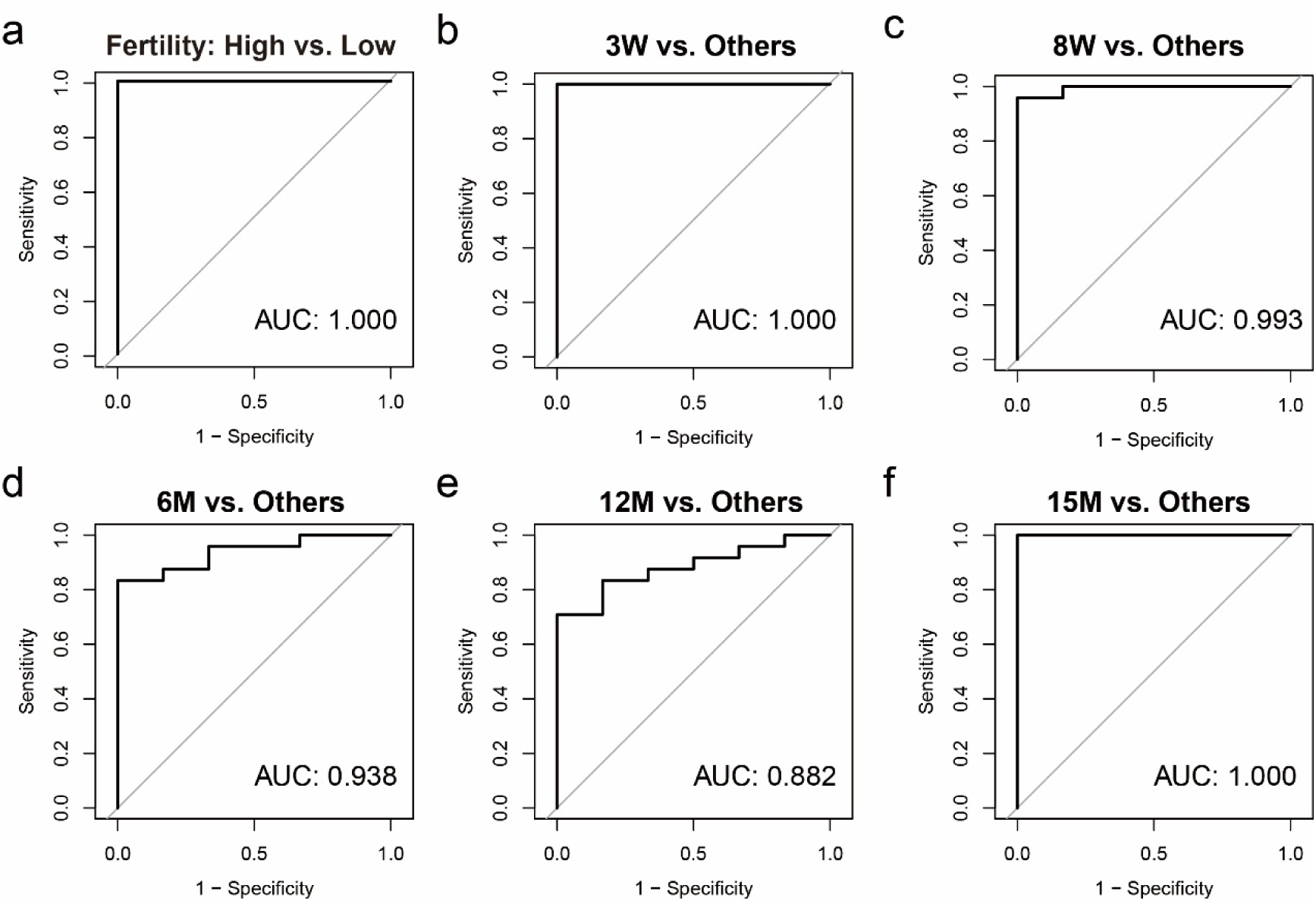
Classification of mouse oocyte aging. **a** The 10-fold cross validation area under the curve (AUC) value of the trained hybrid support vector machine (SVM)-random forest (RF) model for prediction of high versus low fertility. For training, the high fertility group included of 6M and 8W data, while 3W, 12M and 15M data formed the low fertility group. **b-f** The 10-fold cross validation AUC values of the trained hybrid SVM-RF model for predicting 3W (b), 8W (c), 6M (d), 12M (e) and 15M (f) against the corresponding rest groups, respectively. 3W, 8W, 6M, 12M, and 15M stand for 3-week, 8-week, 6-month, 12-month, and 15-month samples, respectively.

## CONCLUSIONS

In summary, we developed an easy-to-use one-pot method for the pretreatment of oocyte samples. The entire workflow is seamlessly integrated into a single step, enabling high throughput with minimal variation. Conventional “elution-drying-resuspension” procedures before LC-MS injection were omitted in our protocol to avoid sample loss. To ensure high success rates, we replaced widely-used 384-well plates with autosampler vials. Importantly, our method requires neither costly equipment or devices nor special reagents, and it can be easily automated and applied to other large cell types. Using this approach, we quantified over 6,500 protein groups from single mouse oocytes and systematically compared their proteomic profiles from post-weaning to end of fertility. This allowed us to identify unique molecular characteristics of oocytes at each age. Additionally, a classifier consisting of ten proteins demonstrated high discrimination power, accurately classifying all five age groups and corresponding fertility levels across the entire reproductive lifespan. To the best of our knowledge, this represents the largest single-cell proteomic dataset for mouse oocyte aging to date. We anticipate that our strategy will facilitate widespread applications in proteomics-based embryogenesis, reproductive biology, and aging research at the single-cell level.

## Author Contributions

Hui Zhang: Methodology, Investigation, Formal analysis. Hailu Zhang: Methodology, Investigation. Chuanxi Huang: Formal analysis. Qing Zeng: Investigation. Chunyan Tian: Supervision. Fuchu He: Supervision. Yun Yang: Conceptualization, Supervision, Writing – Review & Editing.

## ACKNOWLEDGMENTS

The authors acknowledge the assistance of Dr. Linhai Xie from National Center for Protein Sciences (Beijing) in data analysis. This work was supported by the Ministry of Science and Technology of the People’s Republic of China (Grant No. 2020YFE0202200) and the Pre-study Project of Phronesis Medicine Large-scale Scientific Facility funded by Guangzhou Development District.

## Notes

The authors declare no competing financial interest.

## REFERENCES

1 F. A. Rosenberger, M. Thielert and M. Mann, Nat Methods, 2023, 20, 320–323.

2 Nat Methods, 2023, 20, 317–318.

3 D. P. Nusinow, J. Szpyt, M. Ghandi, C. M. Rose, E. R. McDonald, M. Kalocsay, J. Jané-Valbuena, E. Gelfand, D. K. Schweppe, M. Jedrychowski, J. Golji, D. A. Porter, T. Rejtar, Y. K. Wang, G. V. Kryukov, F. Stegmeier, B. K. Erickson, L. A. Garraway, W. R. Sellers and S. P. Gygi, Cell, 2020, 180, 387–402.e16.

4 H. Zhang, S. Ji, K. Zhang, Y. Chen, J. Ming, F. Kong, L. Wang, S. Wang, Z. Zou, Z. Xiong, K. Xu, Z. Lin, B. Huang, L. Liu, Q. Fan, S. Jin, H. Deng and W. Xie, Genome Biol, 2023, 24, 166.

5 Q. Li, L. Mu, X. Yang, G. Wang, J. Liang, S. Wang, H. Zhang and Z. Li, J Proteome Res.

6 J. Huang, P. Chen, L. Jia, T. Li, X. Yang, Q. Liang, Y. Zeng, J. Liu, T. Wu, W. Hu, K. Kee, H. Zeng, X. Liang and C. Zhou, Advanced Science, DOI:10.1002/advs.202301538.

7 Y. He, H. Yuan, Y. Liang, X. Liu, X. Zhang, Y. Ji, B. Zhao, K. Yang, J. Zhang, S. Zhang, Y. Zhang and L. Zhang, Chem Sci, 2023, 14, 13495–13502.

8 Y.-R. Jiang, L. Zhu, L.-R. Cao, Q. Wu, J.-B. Chen, Y. Wang, J. Wu, T.-Y. Zhang, Z.-L. Wang, Z.-Y. Guan, Q.-Q. Xu, Q.-X. Fan, S.-W. Shi, H.-F. Wang, J.-Z. Pan, X.-D. Fu, Y. Wang and Q. Fang, Cell Rep, 2023, 42, 113455.

9 M. Vander Borght and C. Wyns, Clin Biochem, 2018, 62, 2–10.

10 Q. Sang, P. F. Ray and L. Wang, Science (1979), 2023, 380, 158–163.

11 C. Ctortecka, D. Hartlmayr, A. Seth, S. Mendjan, G. Tourniaire, N. D. Udeshi, S. A. Carr and K. Mechtler, Molecular & Cellular Proteomics, 2023, 22, 100665.

12 J. Ma, T. Chen, S. Wu, C. Yang, M. Bai, K. Shu, K. Li, G. Zhang, Z. Jin, F. He, H. Hermjakob and Y. Zhu, Nucleic Acids Res, 2019, 47, D1211–D1217.

13 L. Gatto, R. Aebersold, J. Cox, V. Demichev, J. Derks, E. Emmott, A. M. Franks, A. R. Ivanov, R. T. Kelly, L. Khoury, A. Leduc, M. J. MacCoss, P. Nemes, D. H. Perlman, A. A. Petelski, C. M. Rose, E. M. Schoof, J. Van Eyk, C. Vanderaa, J. R. Yates and N. Slavov, Nat Methods, 2023, 20, 375–386.

14 Z. Y. Li, M. Huang, X. K. Wang, Y. Zhu, J. S. Li, C. C. L. Wong and Q. Fang, Anal Chem, 2018, 90, 5430–5438.

15 Y. Zhu, G. Clair, W. B. Chrisler, Y. Shen, R. Zhao, A. K. Shukla, R. J. Moore, R. S. Misra, G. S. Pryhuber, R. D. Smith, C. Ansong and R. T. Kelly, Angewandte Chemie - International Edition, 2018, 57, 12370–12374.

16 Y. Zhu, P. D. Piehowski, R. Zhao, J. Chen, Y. Shen, R. J. Moore, A. K. Shukla, V. A. Petyuk, M. Campbell-Thompson, C. E. Mathews, R. D. Smith, W. J. Qian and R. T. Kelly, Nat Commun, 2018, 9, 1–10.

17 Q. Lou, Y. Ma, S.-P. Zhao, G.-S. Du and Q. Fang, Talanta, 2021, 224, 121874.

18 J. Woo, S. M. Williams, L. M. Markillie, S. Feng, C. F. Tsai, V. Aguilera-Vazquez, R. L. Sontag, R. J. Moore, D. Hu, H. S. Mehta, J. Cantlon-Bruce, T. Liu, J. N. Adkins, R. D. Smith, G. C. Clair, L. Pasa-Tolic and Y. Zhu, Nat Commun.

19 Z. Yang, K. Jin, Y. Chen, Q. Liu, H. Chen, S. Hu, Y. Wang, Z. Pan, F. Feng, M. Shi, H. Xie, H. Ma and H. Zhou, JACS Au, DOI:10.1021/jacsau.4c00027.

20 S. Karagach, J. Smollich, O. Atrakchi, V. Mohan and T. Geiger, 2024, preprint.

21 M. Matzinger, E. Müller, G. Dürnberger, P. Pichler and K. Mechtler, Anal Chem, 2023, 95, 4435–4445.

22 J. V. V. Isola, S. R. Ocañas, C. R. Hubbart, S. Ko, S. A. Mondal, J. D. Hense, H. N. C. Carter, A. Schneider, S. Kovats, J. Alberola-Ila, W. M. Freeman and M. B. Stout, Nat Aging, 2024, 4, 145–162.

23 M. Li, C. Ren, S. Zhou, Y. He, Y. Guo, H. Zhang, L. Liu, Q. Cao, C. Wang, J. Huang, Y. Hu, X. Bai, X. Guo, W. Shu and R. Huo, Aging Cell.

24 S. Dutta and P. Sengupta, Life Sci, 2016, 152, 244–248.

25 H. Zhang, Z. Shigang and Z. Han, Chinese Journal of Cell Biology, 2024, 46, 646–656.

26 Z. Ye, P. Sabatier, J. Martin-Gonzalez, A. Eguchi, M. Lechner, O. Østergaard, J. Xie, Y. Guo, L. Schultz, R. Truffer, D. B. Bekker-Jensen, N. Bache and J. V. Olsen, Nat Commun.

27 G. Sahoo, D. Samal, P. Khandayataray and M. K. Murthy, Mol Neurobiol, 2023, 60, 5805–5837.

28 R. P. H. De Maeyer and E. S. Chambers, Immunol Lett, 2021, 230, 1–10.

29 Y. Cao, Y. Fan, F. Li, Y. Hao, Y. Kong, C. Chen, X. Hao, D. Han, G. Li, Z. Wang, C. Song, J. Han and H. Zeng, Immunity & Ageing, 2022, 19, 63.

30 Q. Zeng, Y. Gong, N. Zhu, Y. Shi, C. Zhang and L. Qin, Ageing Res Rev, 2024, 97, 102294.

31 M. A. J. Smits, B. V Schomakers, M. van Weeghel, E. J. M. Wever, R. C. I. Wüst, F. Dijk, G. E. Janssens, M. Goddijn, S. Mastenbroek, R. H. Houtkooper and G. Hamer, Human Reproduction, 2023, 38, 2208–2220.

32 J. van der Reest, G. Nardini Cecchino, M. C. Haigis and P. Kordowitzki, Ageing Res Rev, 2021, 70, 101378.

33 P. May-Panloup, L. Boucret, J.-M. Chao de la Barca, V. Desquiret-Dumas, V. Ferré-L’Hotellier, C. Morinière, P. Descamps, V. Procaccio and P. Reynier, Hum Reprod Update, 2016, 22, 725–743.

34 M. Wilding, B. Dale, M. Marino, L. di Matteo, C. Alviggi, M. L. Pisaturo, L. Lombardi and G. De Placido, Human Reproduction, 2001, 16, 909–917.

35 S. S. Sabharwal and P. T. Schumacker, Nat Rev Cancer, 2014, 14, 709–721.

36 A. Agarwal, A. Aponte-Mellado, B. J. Premkumar, A. Shaman and S. Gupta, Reproductive Biology and Endocrinology, 2012, 10, 49.

37 H. Sasaki, T. Hamatani, S. Kamijo, M. Iwai, M. Kobanawa, S. Ogawa, K. Miyado and M. Tanaka, Front Endocrinol (Lausanne).

38 C. So, K. Menelaou, J. Uraji, K. Harasimov, A. M. Steyer, K. B. Seres, J. Bucevičius, G. Lukinavičius, W. Möbius, C. Sibold, A. Tandler-Schneider, H. Eckel, R. Moltrecht, M. Blayney, K. Elder and M. Schuh, Science (1979).

39 S. Wang, Y. Zheng, J. Li, Y. Yu, W. Zhang, M. Song, Z. Liu, Z. Min, H. Hu, Y. Jing, X. He, L. Sun, L. Ma, C. R. Esteban, P. Chan, J. Qiao, Q. Zhou, J. C. Izpisua Belmonte, J. Qu, F. Tang and G.-H. Liu, Cell, 2020, 180, 585–600.e19.

