## Supplementary Material for "Simple One-step Vial-based Pretreatment for Deep Single-cell Proteomics and Its Application to Oocyte Aging"

### Contents

**NanoLC-MS/MS analysis on timsTOF SCP** S3

**Data analysis** S3

**Bioinformatics and statistical analysis** S4

**Fertility classification by machine learning** S5

**Figure S1.** S6

**Figure S2.** S7

**Figure S3.** S8

**Figure S4.** S9

**References** S10

#### ■ EXPERIMENTAL SECTION

**NanoLC-MS/MS analysis on timsTOF SCP**

Nano liquid chromatography-tandem mass spectrometry (nanoLC-MS/MS) analysis was conducted employing a hybrid trapped ion mobility spectrometry (TIMS) quadrupole time-of-flight mass spectrometer (timsTOF SCP, Bruker Daltonics) along with a CaptiveSpray nano-electrospray ion source. Peptide samples underwent separation using a nanoElute ultra-high-performance liquid chromatography (UHPLC) system connected online to the timsTOF SCP. This system employed integrated spraytip columns (50 μm i.d. x 20 cm) packed with 1.9 μm/120 Å ReproSil-Pur C18 resins (Dr. Maisch GmbH, Germany). The mobile phases, referred to as solvents A and B, consisted of water and acetonitrile with 0.1% formic acid (FA), respectively. The flow rate during gradient elution was maintained at 100 nL/min, but was increased to 200 nL/min for the initial 5 minutes. Peptide separation was accomplished over a 60-min total gradient, transitioning from 4% to 85% buffer B (v/v) in the following stages: 4%–6% (v/v) buffer B over 5 minutes, 6%–7% (v/v) buffer B over 3 minutes, 7%–20% (v/v) buffer B over 34 minutes, and 20%–37% (v/v) buffer B over 6 minutes. Columns were heated at 60°C, and each sample was spiked with 0.4 μL of 1x iRT kit (Biognosys) to facilitate retention time calibration.

Mass spectrometry (MS) analysis was performed in diaPASEF mode, scanning precursors across an m/z range of 100–1,700. Precursors with m/z 359 to 959 were selected for fragmentation in 9 PASEF scans, each encompassing 2 ion mobility steps. An isolation window of 25 Th was restricted for each step, with 1 Th overlapping between adjacent windows. Ion mobility resolution was constrained between 1.60 Vs $cm^{-2}$ and 0.57 Vs $cm^{-2}$. The accumulation time and ramp time were 166 ms. Notably, during PASEF MS/MS acquisition, collision energy transitioned linearly from 59 eV at 1/K0 = 1.6 Vs $cm^{-2}$ to 20 eV at 1/K0 = 0.6 Vs $cm^{-2}$.

**Data analysis**

RAW data from method development were analyzed using Spectronaut 17 (Biognosys) software in directDIA mode. These data were searched against the human UniProt FASTA database (20,375 entries, downloaded on December 3, 2021) for 293T cell lysate samples, and the mouse protein reference database (17,090 entries, downloaded on December 3, 2021) for mouse oocytes. Cysteine carbamidomethylation was set as a fixed modification, while protein N-terminal acetylation and methionine oxidation were specified as variable modifications. Enzyme specificity was set to trypsin, allowing for a maximum of two missed cleavages. Precursor filtering was set to perform based on Q-values. Quantification results were output with protein and peptide false discovery rates (FDRs) less than 0.01. All other settings were left at their default values.

For mouse oocytes across five age groups, Spectronaut 19 (Biognosys) software was applied instead. Cysteine carbamidomethylation was removed from fixed modifications, cross-run normalization was enabled, while all the other settings were identical to those used for RAW data in method development part.

**Bioinformatics and statistical analysis**

Proteins detected in less than 50% of all samples were excluded. Protein abundance values were normalized using median normalization, and missing values were imputed using the K-Nearest Neighbours (KNN) algorithm. The resulting quantification matrix was then $\log_2$-transformed. Two-tailed unpaired Student's t test or one-way ANOVA was employed for differential expression analysis. Proteins with fold change >1.5 and p-value <0.01 (for ANOVA, significant proteins were identified with a q-value < 0.05) were designated as differentially expressed proteins (DEPs). Data analysis and visualization were performed using custom R (4.2.2) scripts. The following R packages were used: pheatmap (v1.0.12), ggplot2 (v3.4.4), pROC (v1.18.5), and clusterProfiler (v4.2.2). Gene Ontology (GO) enrichment analysis was carried out using the msigdbr (v7.5.1) and clusterProfiler (v4.2.2) packages, with a q-value threshold of 0.05 for significance. Gene expression clustering was performed with the Mfuzz (v2.54.0) package. Statistical analyses and plots for the five reported markers were created using GraphPad Prism (v8).

**Fertility classification by machine learning**

Differential statistical analysis was performed using analysis of variance (ANOVA) tests across groups, and the top 100 DEPs were selected based on adjusted p-values. Next, samples were categorized into two fertility levels: the high fertility group (6-month and 8-month samples) and the low fertility group (remaining samples). Subsequently, using the identified top 100 DEPs, classifier combinations were generated. The dataset was randomly split into a training set (70%, 21 samples) and a testing set (30%, 9 samples). Hybrid classification models were trained using support vector machine (SVM) and random forest (RF) algorithms. The combination of classifiers yielding the highest average F1-score (a weighted average of precision and recall)[1] across both models on the test set was identified. Random forest models were built with 500 trees using the R package randomForest, employing 10-fold cross-validation. Then, the performance of the generated fertility classifier across multiple sets of models was evaluated using leave-one-out cross-validation (LOOCV). Prediction performance metrics, including the area under the curve (AUC) values, were calculated for all samples.

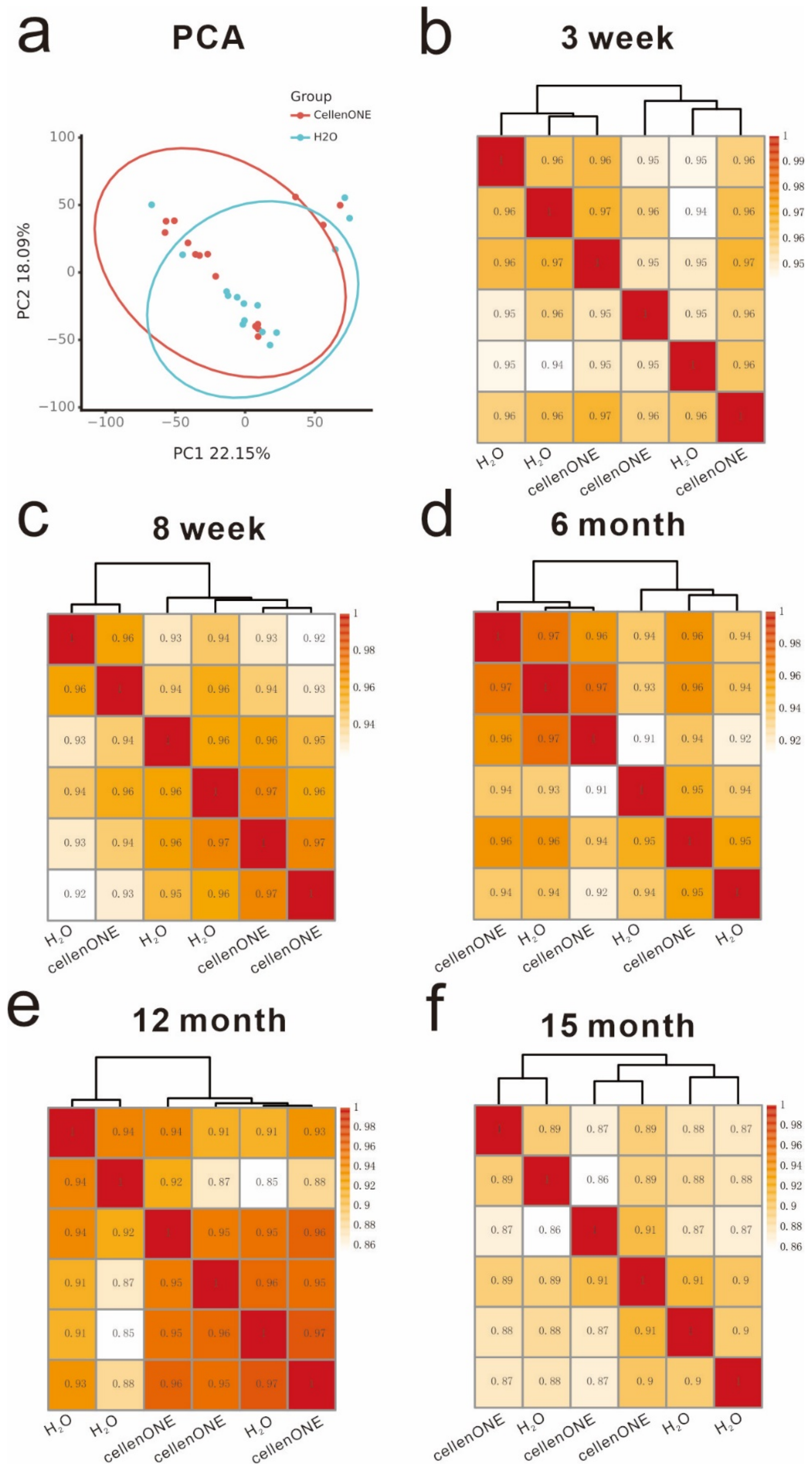


**Figure S1.** PCA analysis of all data (a) and heatmaps depicting the Pearson correlation coefficients of the proteome across biological replicates in five age groups: 3 week (b), 8 week (c), 6 month (d), 12 month (e) and 15 month (f).

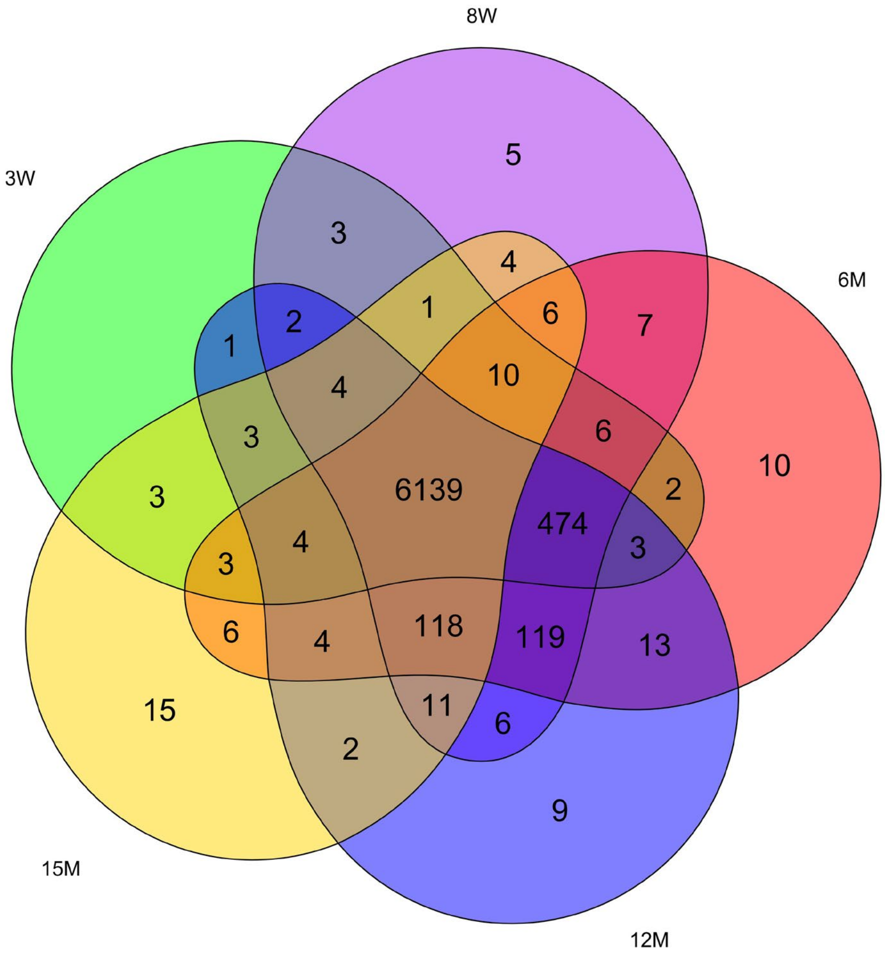


**Figure S2.** Venn diagrams depicting quantified protein groups across the five age groups.

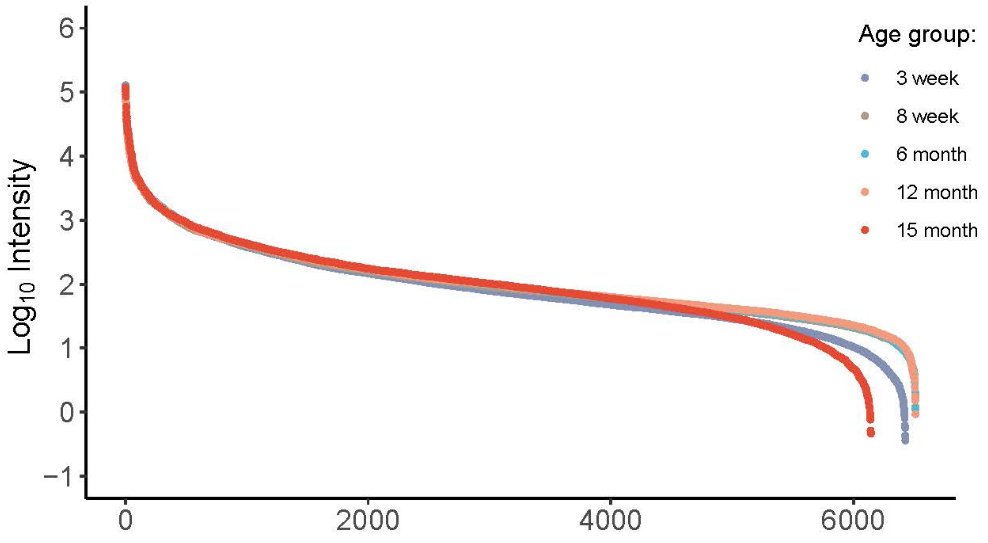


**Figure S3**. Dynamic ranges of quantified protein groups across the five age groups.

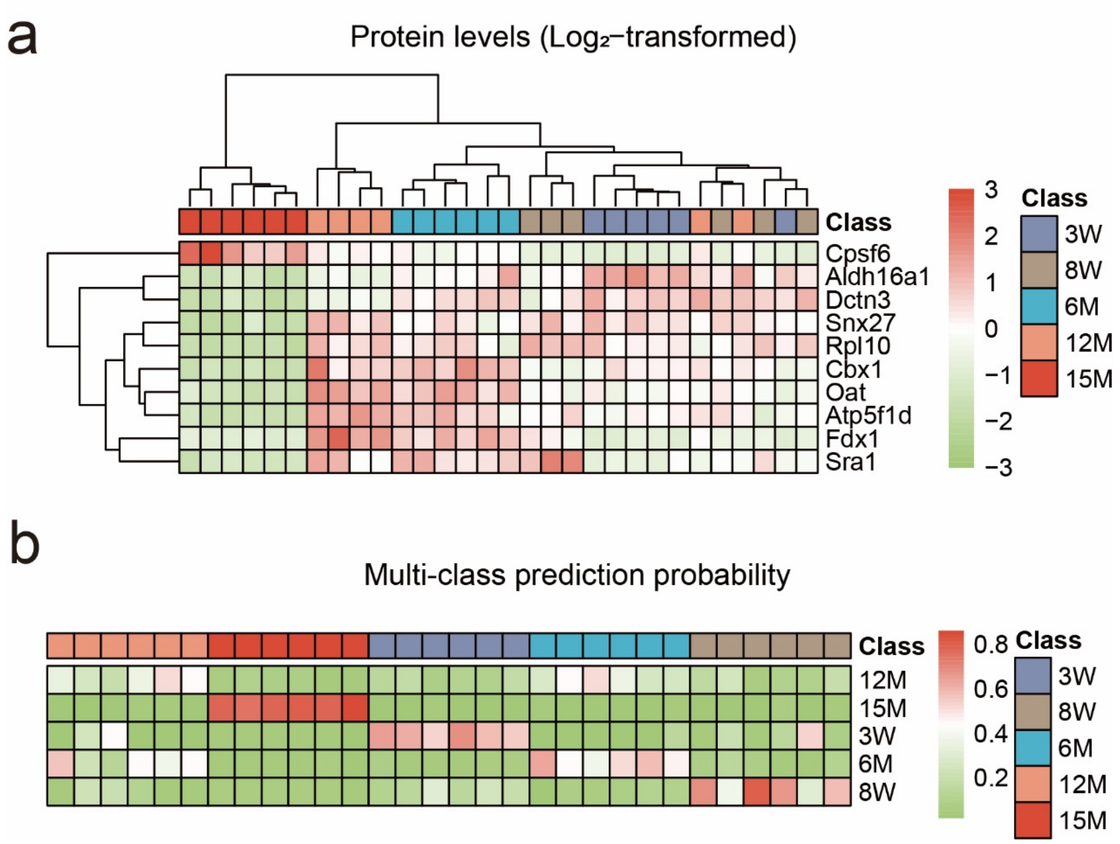


**Figure S4**. Protein levels (a) and prediction probabilities (b) for the 10 proteins included in the classifier across the five age groups.

**REFERENCES**

1 W. Tun, J. K.-W. Wong and S.-H. Ling, *Sensors*, 2021, 21, 8163.
